# Neurotransmitter Genes in the Nucleus Accumbens that Are Involved in the Development of Behavioral Pathology After Positive Fighting Experiences and Their Deprivation. A Conceptual Paradigm for Neurogenomic Data Analysis

**DOI:** 10.1101/2023.12.30.573683

**Authors:** Natalia N. Kudryavtseva, Dmitry A. Smagin, Olga E. Redina, Irina L. Kovalenko, Anna G. Galyamina, Vladimir N. Babenko

**Affiliations:** Neuropathology Modeling Laboratory, Neurogenetics of Social Behavior Sector, FRC Institute of Cytology and Genetics, SB RAS, Novosibirsk, Russia; (N.N.K); (D.A.S.); (O.E.R.); (I.L.K.); (A.G.G.); (V.N.B.)

**Keywords:** nucleus accumbens, positive fighting experience, fighting deprivation, neurotransmitter differentially expressed gene

## Abstract

It has been shown earlier that repeated positive fighting experience in daily agonistic interactions is accompanied by the development of psychosis-like behavior with signs of an addiction-like state associated with changes in the expression of genes encoding the proteins involved in the main neurotransmitter events in some brain regions of aggressive male mice. Fighting deprivation (a no-fight period of 2 weeks) causes a significant increase in their aggressiveness. This paper is aimed at studying—after a period of fighting deprivation—the involvement of genes (associated with neurotransmitter systems within the nucleus accumbens) in the above phenomena. The nucleus accumbens is known to participate in reward-related mechanisms of aggression. We found the following differentially expressed genes (DEGs), whose expression significantly differed from that in controls and/or mice with positive fighting experience in daily agonistic interactions followed by fighting deprivation: catecholaminergic genes *Th*, *Drd1*, *Drd2*, *Adra2c*, *Ppp1r1b*, and *Maoa*; serotonergic genes *Maoa*, *Htr1a*, *Htr1f*, and *Htr3a*; opioidergic genes *Oprk1*, *Pdyn*, and *Penk*; and glutamatergic genes *Grid1*, *Grik4*, *Grik5*, *Grin3a*, *Grm2*, *Grm5*, *Grm7*, and *Gad1.* The expression of DEGs encoding proteins of the GABAergic system in experienced aggressive male mice mostly returned to control levels after fighting deprivation except for *Gabra5*. In light of the conceptual paradigm for analyzing data that was chosen in our study, the aforementioned DEGs associated with the behavioral pathology can be considered responsible for consequences of aggression followed by fighting deprivation, including mechanisms of an aggression relapse.

## 1. Introduction

Effects of repeated experience of own aggression on behavior, physiology, and neurochemistry in animals have been studied purposefully by J.P. Scott [1,2], P.F. Brain with colleagues [3–7], and J.M. Koolhaas with colleagues [8,9]. Research on the influence of positive fighting experience in daily agonistic interactions on mice has also been the focus of N.N. Kudryavtseva with colleagues for a long period [e.g., 10-24]. Experimental data indicate that male mice that had a repeated aggressive experience accompanied by wins develop a behavioral psychopathology [11–13,17,18], which has been verified by data on the development of psychoneurological symptoms. It has been shown that repeatedly aggressive and winning male mice after 20 days of agonistic interactions demonstrate changes in individual and social behaviors such as enhanced impulsivity and aggressiveness, disturbances in social recognition, hyperactivity, stereotypic uncontrolled behaviors, and symptoms of autistic spectrum disorders. After a period of fighting deprivation (without agonistic interactions for 2 weeks), these male mice demonstrate increased aggression (in comparison with the period before the deprivation), which is accompanied by preservation or enhancement of all types of pathological behavior seen in the aggressive animals [13,25,26]. It has been obvious that the repeated positive fighting experience in daily agonistic interactions with mouse male partners is accompanied by the development of psychosis-like behavior with signs of an addiction-like state [12,13,25,26] or manic-like behavior [27] in mice. A neurochemical study on brain changes in aggressive male mice has revealed activation of dopaminergic systems in the ventral tegmental area (VTA) and in the dorsal striatum (STR) and inhibition of serotonergic-system activity in midbrain raphe nuclei (MRNs) [11–13].

It has been well known that the main neurotransmitter systems—catecholaminergic (CAergic), serotonergic, opioidergic, GABAergic, and glutamatergic—are involved in the mechanisms of reward, drug abuse, addiction, and aggression [e.g., 6,12,28-33]. Our neurotranscriptomic and cellular studies on mice experienced in aggression have revealed alterations of the expression of numerous genes specific for brain regions depending on their main functions in the regulation of aggression mechanisms [16,18–24,34]. For example, in the VTA, which is the main brain region involved in reward mechanisms, an increase has been revealed in the expression of CAergic (*Th*, *Ddc*, *Slc6a2*, and *Slc6a3*), glutamatergic (*Slc17a7* and *Slc17a8*), and opioidergic (*Oprk1)* genes (and others) encoding relevant proteins [16]. Moreover, we have previously shown that in chronically winning (aggressive) mice, there is an increase in the proliferation of progenitor neurons and in the production of young neurons in the dentate gyrus (in the hippocampus), and these neurogenesis parameters remain modified during 2 weeks of fighting deprivation [15]. This means that hippocampal transcriptomic changes are associated with enhanced neurogenesis and take part in the formation of behavioral features in mice having the positive fighting experience in agonistic interactions as well as during the subsequent fighting deprivation. Some of these genes in the hippocampus have been associated with behavioral traits, including abnormal aggression-related behavior, an abnormal anxiety-related response, and other traits [24]. The transcriptomic changes of gene expression get reversed only partially after a period of no fights. Similar data have been obtained about the STR [21].

We believe that genes encoding proteins responsible for the aggravation of the pathological condition in aggressive mice after 2-week deprivation of fighting (aggression relapse) retain the altered expression (as compared to the control level) after the no-fight period. This supposition is based on our observations of alterations of social and individual behaviors and of the emotional state [reviews: 12,13,17,25] that persist in aggressive mice for at least 2 weeks after the removal of the psychopathogenic external factor, that is, after a period without agonistic interactions; this persistence can serve as an indicator of the development of a psychoneurological disorder. Here, we are extending this view to other psychopathologies that may arise in mice in the course of a similar experiment. Our conceptual paradigm of data analysis is based on the notion that a disease manifestation can result from the participation of pathology-related genes that retain altered expression for a long time even after cessation of the exposure to a pathological stimulus.

There has been interest in the role of the mesolimbic dopaminergic circuit in the control over aggression reward [review 35-37] because dopaminergic projections from the VTA to the nucleus accumbens (NAcc) modulate both aggression intensity and dopamine (DA) levels [38,39]. Our attention in this research has been concentrated on neurotransmitter genes in the NAcc, which is involved in reward mechanisms [36,40,41] and in consequences of the repeated experience of aggression in mice [16,19,21,24].

The present study is focused on the differentially expressed genes (DEGs) associated with CAergic, serotonergic, opioidergic, glutamatergic, and GABAergic systems’ activities in the NAcc of mice having the repeated positive aggressive experience for 20 days and those after 2-week deprivation of fighting. To obtain these data, transcriptomic analysis was performed. The study was carried out within the research field that we are developing, “Functional neurotranscriptomics of pathological conditions,” by means of the sensory contact model [10], which was later renamed as the “chronic social conflict model.” In this experiment, three groups of animals were analyzed: (1) controls, i.e., mice without a consecutive experience of agonistic interactions (C); (2) A20 winners, i.e., a group of mice aggressive daily for 20 days; and (3) AD (aggression deprivation), i.e., fight-deprived winners that were repeatedly aggressive for 20 days and then were subjected to 14-day deprivation (the no-fight period, without agonistic interactions). A graphical outline of the experiment is given in the section Materials and Methods.

## 2. Results

### 2.1. The Winners’ Behavior During Agonistic Interactions

Among the winners with a positive fighting experience in 20 tests (for 20 days), the most aggressive animals were identified, which demonstrated in daily agonistic interactions the longest time of aggressive motivation (in seconds) as a behavioral reaction to a partner in the neighboring compartment of the cage before agonistic interaction in the partition test, and hostile behaviors in the agonistic interaction test including the amount of time (s): total duration of attacks, duration (s) of aggressive grooming and diggings (see a detailed description in the Materials and Methods section, Figure 1).

**Figure 1.**
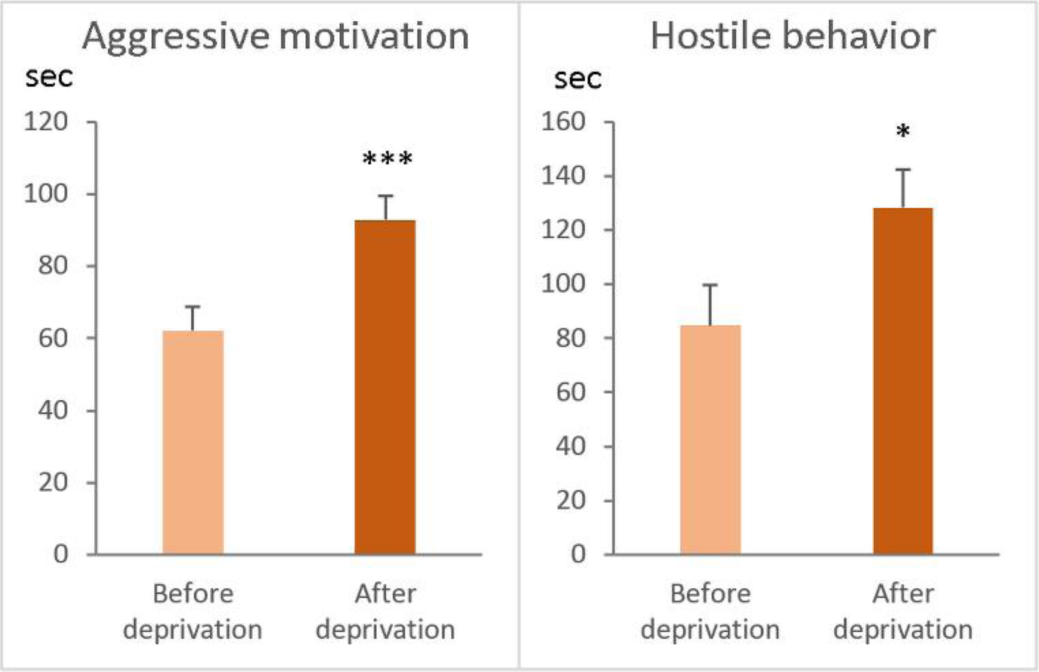
Total duration of aggressive motivation and hostile behavior demonstrated by experienced aggressive mice before (A20 mice, n = 10) and after fighting deprivation (AD mice, n = 10). \**p* < 0.05; \*\*\**p* < 0.001, Wilcoxon test.

Using the timetable of the experiment, it was shown (Figure 1) that aggressive motivation (total time spent near the partition as a reaction to the partner in the neighboring compartment of the cage) and total duration of hostile behavior during the agonistic test were significantly greater in AD mice (after deprivation) than in A20 mice (before deprivation). This means that experimental groups of mice demonstrated the main symptom of the behavioral pathology: an increase in aggressiveness after the period of deprivation as well as enhanced hyperactivity and aggressive motivation toward unaggressive partners.

### 2.2. Detecting DEGs in Pairwise Comparisons

Comparisons of gene expression data between groups control, A20, and AD groups were performed according to ref. [42]. This analysis yielded approximately 2629 DEGs for the “control vs A20” comparison, 1111 DEGs for the “control vs. AD” comparison (Table 1), and 2578 DEGs for the “A20 vs. AD” comparison. This finding indicated that in NAcc of AD mice, the number of DEGs is decreased, tending to return to the control level.

**Table 1.** Three-way comparisons based on detection of DEGs.

| Number of DEGs (FDR < 0.05) |  |
| --- | --- |
| Control vs. A20 | 2629 |
| Control vs. AD | 1111 |
| A20 vs. AD | 2578 |

It was found (Figure 2, Supplementary Table S1) that four out of 14 DEGs in the control vs. A20 comparison were upregulated: *Th*, *Slc6a3*, *Adra1a*, and *Slc18a2*. Genes *Adra2a*, *Drd1*, *Maoa*, *Maob*, *Ppp1r1b*, and *Snca* were downregulated. After the period of aggression deprivation (control vs. AD comparison), a return to the control level of expression was registered for genes *Slc6a3*, *Adra1a*, *Adra2a*, *Maob*, *Snca*, and *Slc18a2*. In this comparison, there was increased expression of genes *Adra2c*, *Drd1*, *Drd2*, and *Ppp1r1b* and decreased expression of the *Th* gene in AD mice) relative to the control and to A20 mice. The expression of *Drd3* and *Adrb1* in AD mice did not differ from that in the control but was higher than that in A20 mice.

**Figure 2.**
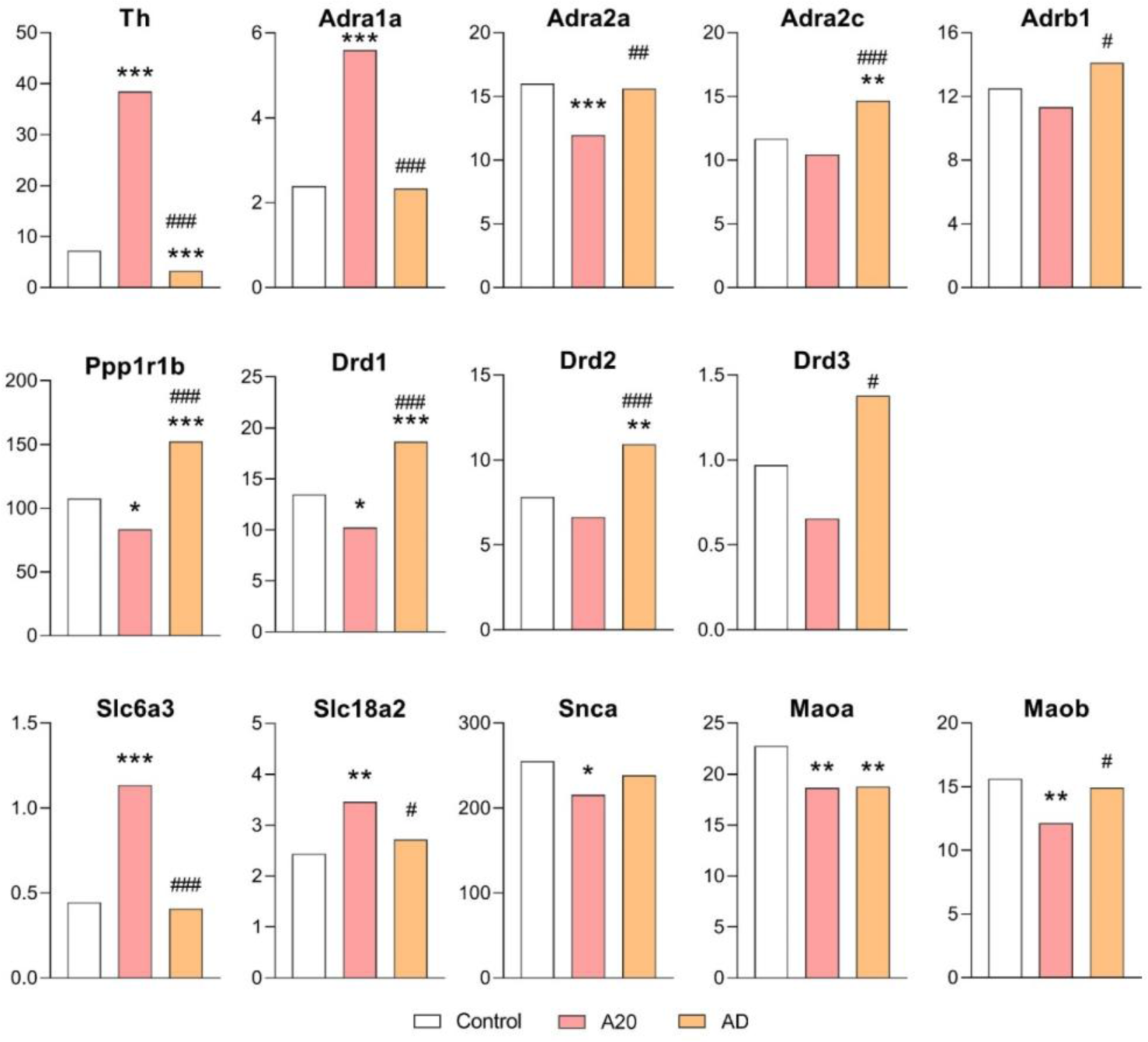
CAergic-system–associated DEGs in the NAcc. *Control (C) vs. A20 and C vs. AD; ^#^A20 vs. AD. Data are presented in FPKM units (fragments per kilobase of transcript per million mapped reads); C: male mice with no consecutive experience of agonistic interactions; A20: aggressive male mice with consecutive 20 days of wins in daily agonistic interactions; AD: A20 males after a period of fighting deprivation for 14 days. \**p* < 0.05; \*\**p* < 0.01; \*\*\**p* < 0.001; ^#^*p* < 0.05; ^##^*p* < 0.01; ^###^*p* < 0.001. Additional information is shown in Supplementary Table S1.

In light of our conceptual paradigm of data analysis, these results of the comparison of control vs. AD mice can be interpreted as follows: *Adra2c* (gene encoding adrenoreceptors 2c), *Drd1* and *Drd2* (genes encoding DA receptors), *Th* (gene encoding the rate-limiting enzyme of the catecholamine synthesis), and *Maoa* (gene coding for mitochondrial enzymes that catalyze oxidative deamination of all monoamines) can be considered genes associated with the pathology. The *Ppp1r1b* gene (encoding the DARPP-32 protein) mediates the actions of DA and regulates the stimulation of dopaminergic and glutamatergic receptors; *Ppp1r1b* shows the highest expression in AD mice in the absence of DA after aggression deprivation [Figure 2]. It is well known that DARPP-32 is crucial for the regulation of transcriptional and behavioral responses to pharmacological agents, including neuroleptics, antidepressants, and drugs of abuse and may serve as a therapeutic target in neurological and psychiatric disorders [43].

We assume that these genes associated with CAergic systems in the NAcc can be responsible for the development of reward-related effects of wins in the agonistic interactions, and these effects are the main symptoms of the pathological state similar to a psychosis-like disorder.

The present neurotranscriptomic analysis confirmed our previous finding that repeated aggression is accompanied by an increase in the activity of brain dopaminergic systems [44] as evidenced by elevated DOPAC (3,4-dihydroxyphenylacetic acid) levels and/or increased DOPAC/DA ratios in the olfactory bulbs, amygdala, hippocampus, NAcc, STR, and midbrain in the winners. Other authors have reported elevation of DA levels in the NAcc before, during, and after fights in aggressive rats [28, 38]. In addition to these data, in the VTA (containing somata of mesolimbic dopaminergic neurons and playing an important role in the mediation of reward processes), overexpression of genes *Th*, *Dat1 (Slc6a3)*, and *Snca* has been found in aggressive winners [34,45]. Elevated mRNA levels of *Th* and *Slc6a3* persisted in the winners after the deprivation period in our work. The persistence of the altered dopaminergic-gene expression confirmed the development of a psychopathology in the winners. Of note, the level of aggressiveness—measured as latency to the first attack, the number of attacks, and total time spent attacking— is reported to positively correlate with *Th* and *Snca* mRNA levels in the VTA [34,45]. Thus, activation of dopaminergic systems during repeated positive fighting experience enhances the expression of dopaminergic genes in the VTA and according to our present experiments in the NAcc, which are the main reward-mediating brain regions.

In our previous study, we used an experimental approach to investigation of a psychotropic drug’s effects under simulated clinical conditions [46]. The pharmacological data confirmed alterations of dopaminergic activity in the brain of repeatedly aggressive mice. Haloperidol, an antagonist of dopaminergic receptors, effectively inhibits aggression in male mice having a short (2-day) experience of aggression but is ineffective in the winners with a 20-day experience [47]. Dopamine D1 receptor antagonist SCH-23390 has a similar effect [48]. This means that dopaminergic receptors may be sensitized or desensitized to the blocking effects of DA receptor antagonists depending on the duration of the aggressive experience.

#### 2.2.1. Opioidergic and cannabinoidergic genes

Increased expression of *Oprd1* gene (encoding opioidergic 1d receptors) and decreased expression of *Cnr1* (encoding cannabinoid Receptor 1, associated with cannabis dependence and abuse) and of *Faah* (fatty acid amide hydrolase, which catalyzes the hydrolysis of the endocannabinoid 2-arachidonoylglycerol) were found in comparison with the control (С vs. A20 comparison; Supplementary Table S1). Nonetheless, their expression returned to the control level after the fighting deprivation (in AD mice).

In the C vs. A20 comparison, there was no significant difference in expression for the following genes: *Oprk1* (encoding kappa opioid receptors) and *Pdyn* (whose product is proteolytically processed to form secreted opioid peptides beta-neoendorphin, dynorphin, leu-enkephalin, rimorphin, and leumorphin: these peptides are ligands for the kappa-type of opioid receptor; dynorphin is involved in modulation of responses to several psychoactive substances, including cocaine). Similar absence of differences in expression was observed for the *Penk* (proenkephalin) gene, which produces pentapeptide opioids Met-enkephalin and Leu-enkephalin, which are stored in synaptic vesicles and then released into the synapse, where they bind to mu- and delta-opioid receptors.

By contrast, after the period of fighting deprivation, the expression of *Oprk1* and *Pdyn* increased significantly in the comparison of A20 vs. AD mice, and *Penk* expression increased significantly in the C vs. AD comparison. Thus, these three genes may participate in the consequences of deprivation and may be involved in the manifestation of an aggression relapse. Additionally, in our previous study, we detected upregulation of opioidergic genes in some brain regions: the *Oprk1* gene in the VTA and *Oprd1* and *Penk* in the prefrontal cortex of A20 mice compared with the control [16].

On the basis of previous pharmacological experiments, we have hypothesized that chronic activation of the brain’s opioidergic systems in aggressive winners leads to opioid drug tolerance similar to that of human addicts. Various experiments have shown that naltrexone (an opioid receptor antagonist) effectively reduces aggressiveness in male mice with a short experience of aggression, and at the same doses, is ineffective in male mice with a long history of aggression [49–51]. Selective mu-opioid receptor agonist morphine has a stimulatory effect on locomotor activity of control mice and has no effect on locomotor activity in 60% of the winners after fighting deprivation [25]. Desensitization of kappa-opioid receptors to selective agonist U50,488 in experienced aggressive winners has been documented too [52,53]. This effect is associated with underexpression of the gene coding for kappa-opioid receptors in the VTA of male mice after 10 days of agonistic interactions [54].

Thus, opioidergic receptors could be sensitized or desensitized depending on the amount of aggressive experience and severity of aggressiveness in mice. In this context, recurrent aggression and its enhancement after fighting deprivation has been explained by a deprivation (withdrawal) effect due to the involvement of opioidergic systems in reward-related effects of repeated aggression. Possibly, similarly to humans, these findings indicate the development of tolerance to endogenous opioids (for example, the brain’s endogenous morphine) [55–57] or an opioid deficiency in the absence of DA inducing positive reinforcement.

It is well known that the opioidergic and cannabinoidergic genes are involved in opioid-mediated mechanisms underlying social interaction, attachment, and aggression [6,58] and in the formation of addiction states during repeated activation by drugs with rewarding effects (inducing good mood and positive emotions in humans). Our current experiments revealed that the acquisition of a positive fighting experience in daily agonistic interactions is accompanied by positive emotions induced by the wins: aggressive males every day demonstrate strong aggressive motivation toward the defeated male mice (losers) and repeatedly attack them even in the absence of the need to demonstrate superiority.

#### 2.2.2. Serotonergic Genes

In this experiment, in six out of the eight genes encoding proteins participating in regulation of serotonergic system activity (*Htr1d*, *Htr1f*, *Htr2c*, *Maoa*, *Maob*, and *Slc18a2*), expression changed in the C vs. A20 comparison (Figures 2 and 4): upregulation of *Slc18a2* and *Htr1f* and downregulation of *Htr2c*, *Htr1d*, *Maoa*, and *Maob* genes. After the period of aggression deprivation (C vs. AD), a return to the control level of expression of the *Htr2c*, *Htr1d*, *Maob*, and *Slc18a2* genes was noted. In this comparison, increased expression of the *Htr1a* gene and decreased expression of *Maoa* gene were registered after the deprivation as compared with the control. Decreased expression of genes *Мaoa*, *Htr1f*, and *Htr3a* and increased expression of *Htr1a*, *Htr1d*, *Htr2c*, and *Maob* were detected in the A20 vs. AD comparison (Figures 2 and 4). We suppose that downregulation of *Htr1f*, *Htr3a*, *Maoa* and upregulation of *Htr1a* genes in comparisons “C vs. A20” and “C vs. AD” may be a specific effect of fighting deprivation, and these genes may be regarded as participating in the development of the pathological state of the AD mice.

**Figure 3.**
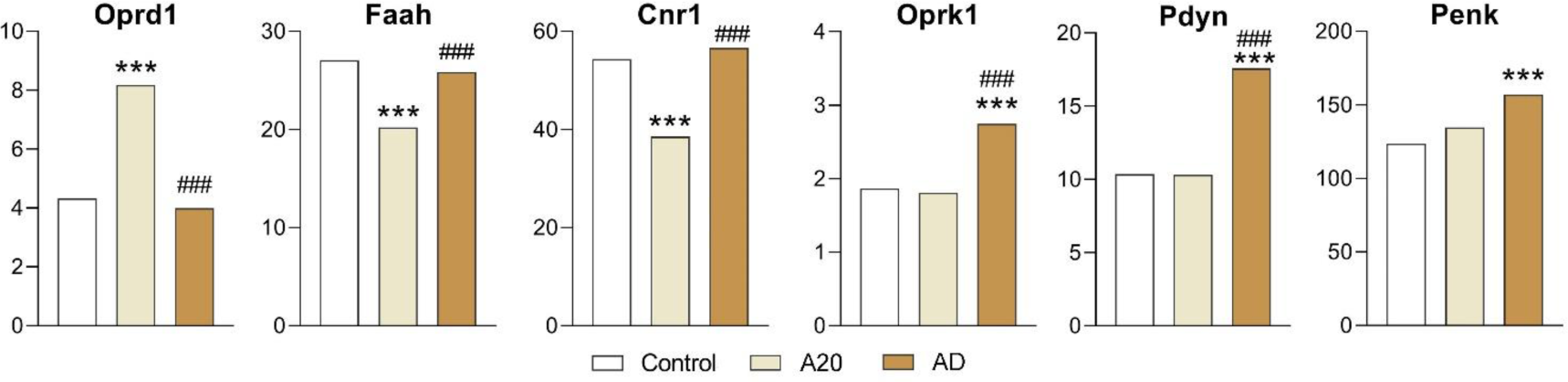
Opioidergic-systemand cannabinoidergic-system–associated DEGs in the NAcc. *Control (C) vs. A20 or C vs. AD; ^#^A20 vs. AD. Data are presented in FPKM units (fragments per kilobase of transcript per million mapped reads); C: male mice without experience of agonistic interactions; A20: aggressive males with consecutive 20 days of wins in daily agonistic interactions; AD: A20 mice deprived of fighting for 14 days. \*\*\**p* < 0.001; ^###^*p* < 0.001. Additional information is shown in Supplementary Table S1.

**Figure 4.**
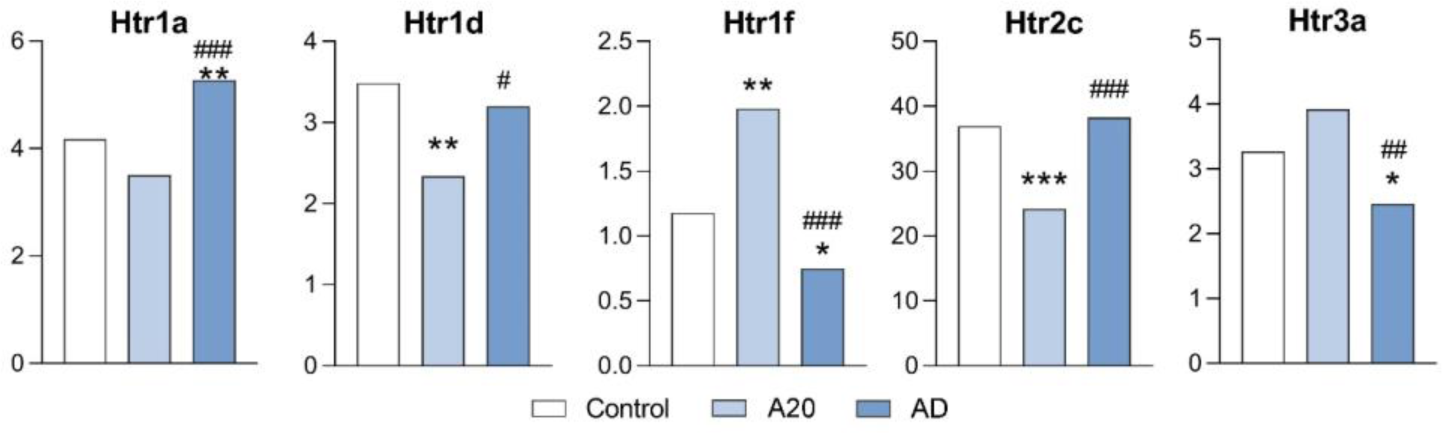
Serotonergic-system–associated DEGs in the NAcc. *Control (C) vs. A20 or C vs. AD; ^#^A20 vs. AD. Data are presented in FPKM units (fragments per kilobase of transcript per million mapped reads); C: male mice with no experience of agonistic interactions; A20: males with consecutive 20 days of wins in daily agonistic interactions; AD: males with consecutive 20 days of wins next deprived of fighting for 14 days. \**p* < 0.05; \*\**p* < 0.01; \*\*\**p* < 0.001; ^#^*p* < 0.05; ^##^*p* < 0.01; ^###^*p* < 0.001. Additional information is given in Supplementary Table S1.

A reduced 5-hydroxyindoleacetic acid (5-HIAA) level and/or 5-HIAA/5-HT ratio [11,12,44] and a decrease in the activity of a rate-limiting enzyme of serotonin (5-HT) biosynthesis (i.e., tryptophan hydroxylase) in the midbrain and STR have been demonstrated earlier in male mice with repeated aggressive experience [59,60]. Later, a transcriptomic analysis has revealed alterations of the expression of serotonergic genes involved in the synthesis, inactivation, receptor sensitivity, as detected in different brain regions; for example, in aggressive mice with repeated positive fighting experience, genes *Tph2*, *Ddc*, *Slc6a4*, *Htr2a*, *Htr3a*, and *Htr5b* are downregulated in MRNs, containing mostly the somata of serotonergic neurons [14,61]. In the hypothalamus, genes *Maoa*, *Htr2a*, *Htr2c* are downregulated too. These data have confirmed inhibition of serotonergic activity under the influence of repeated aggression [11]. By contrast, after the period of fighting deprivation, the expression of the *Tph2* and *Slc6a4* (Sert) genes is significantly higher as compared to the period before the deprivation, thereby returning to the normal (control) level, while the mRNA level of *Maoa* and *Htr1a* remains at a significantly higher level in MRNs as compared with respective controls [14]. The change in serotonergic activity during the acquisition of aggressive experience has also been confirmed by pharmacological studies. Specific changes in pharmacological sensitivity of 5-HT1A receptors to agonist buspirone has been documented [62]. All these findings are consistent with the hypothesis of 5-HT deficiency in the brain of mice experienced in aggression [11,12].

The concept of a major inhibitory influence of 5-HT on aggressive behavior in animals and humans has been widely accepted by many investigators. Reduced cerebrospinal-fluid levels of a major 5-HT metabolite called 5-HIAA as well as decreased sensitivity of 5-HT1A receptors have been shown in prisoners or psychiatric patients with a history of repeated violent or aggressive acts [63–65] or in rodents and nonhuman primates [66], thus possibly indicating a reduced inhibitory serotonergic function in aggressive individuals.

We propose that the return of the expression of *Htr2c*, *Htr1d*, *Maob*, and *Slc18a2* genes to the control level after the period of fighting deprivation implies that changes in brain serotonergic activity in the NAcc are not the main cause of the behavioral pathology developing in the male mice in this experimental context. It has been suggested that reduced serotonergic activity stimulates the manifestation of aggression in conflict situations owing to the development of increased impulsivity [11,61]. On the contrary, here we show that changes in activity of the serotonergic system in the NAcc may be related to expression levels of genes *Htr1a*, *Htr1f*, *Htr3a*, and *Maoa*. Moreover, the roles of different brain regions in the formation of impulsivity in individuals may differ.

In addition, our data support the idea that the repeated experience of aggression creates an imbalance between activities of two neurotransmitter systems—serotonergic and dopaminergic [12]—namely, inhibition and activation of these monoaminergic systems, respectively.

#### 2.2.3. GABAergic Genes

**Figure 5.**
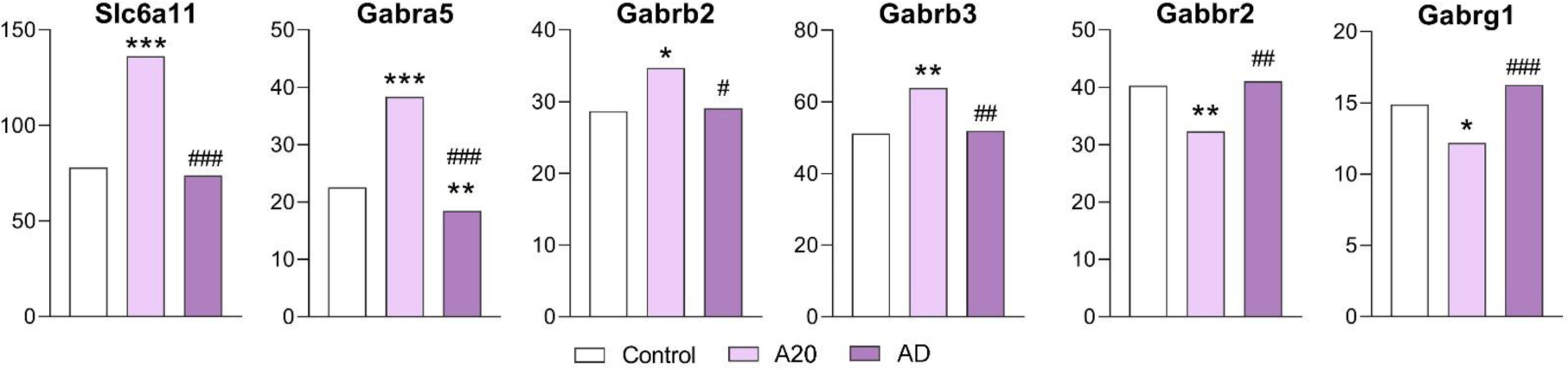
GABAergic-system–associated DEGs in the NAcc. *Control (C) vs. A20 or C vs. AD; ^#^A20 vs. AD. Data are presented in FPKM units (fragments per kilobase of transcript per million mapped reads); C: mice with no experience of agonistic interactions; A20: male mice with consecutive 20 days of wins in daily agonistic interactions; AD: male mice with consecutive 20 days of wins in daily agonistic interactions next deprived of fighting for 14 days. \**p* < 0.05; \*\**p* < 0.01; \*\*\**p* < 0.001; ^#^*p* < 0.05; ^##^*p* < 0.01; ^###^*p* < 0.001. Additional information is presented in Supplementary Table S1.

The expression of only six out of the 23 genes related to the functioning of the GABAergic system was found to change: expression of *Slc6a11*, *Gabra5*, *Gabrb2*, and *Gabrb3* increased, and *Gabbr2* and *Gabrg1* expression decreased in the C vs. A20 comparison. After the period of deprivation, expression of all genes, except for the *Gabra5* gene, returned to the control levels.

Therefore, the GABAergic system in the NAcc can be considered a system participating in the development of the behavioral pathology during repeated aggression but seems not to play a crucial part in the manifestation of the aggression relapse. Of note, earlier we have detected downregulation of GABAergic genes in the STR (*Gabra2*, *Gabra3*, *Gabrb2*, *Gabrg2*, and *Gabrg3)* and upregulation of some of these genes in the VTA (*Gabra1* and *Gabrg2)* of A20 mice [16]. Those data led us to suppose that the changes in gene expression may depend on the neurochemical environment, which differs among brain regions.

#### 2.2.4. The Glutamatergic genes

Among 24 glutamatergic DEGs in A20 and/or AD mice, there were seven genes coding for metabotropic receptors (*Grm1*, *Grm2*, *Grm3*, *Grm4*, *Grm5*, *Grm7*, and *Grm8*), 12 genes encoding ionotropic receptors (*Gria3a Gria4*, *Grid1*, *Grid2ip*, *Grik1*, *Grik3*, *Grik4*, *Grik5*, *Grin1*, *Grin2a*, *Grin2d*, and *Grin3a*), two genes encoding glutamate decarboxylase (*Gad1* and *Gad2*), and two genes coding for inorganic transporters: *Slc17a7* and *Slc17a8* (Supplementary Table S1).

In the comparison of C vs. А20 mice, 13 genes *(Grin2a*, *Grin2d*, *Grid2ip*, *Grik1*, *Grik3*, *Gria4*, *Grin1*, *Grm1*, *Grm2*, *Grm4*, *Grm8*, *Gad1*, and *Gad2)* proved to be upregulated. Four genes *(Gria3*, *Grm3*, *Slc17a7*, and *Slc17a8)* were found to be downregulated. For seven genes *(Grid1*, *Grin3a*, *Grik4*, *Grik5*, *Grm5*, and *Slc17a6)*, expression was unchanged (Supplementary Table S1).

In the comparison of C vs. АD mice, 15 genes *(Gad2*, *Gria3*, *Gria4*, *Grid2ip*, *Grik1 Grik3*, *Grin1*, *Grin2a*, *Grm1*, *Grm3*, *Grm4*, *Grm8*, *Grin2d*, *Slc17a7*, and *Slc17a8)* returned to the control expression level after aggression deprivation (Supplementary Table S1, Figure 6).

**Figure 6.**
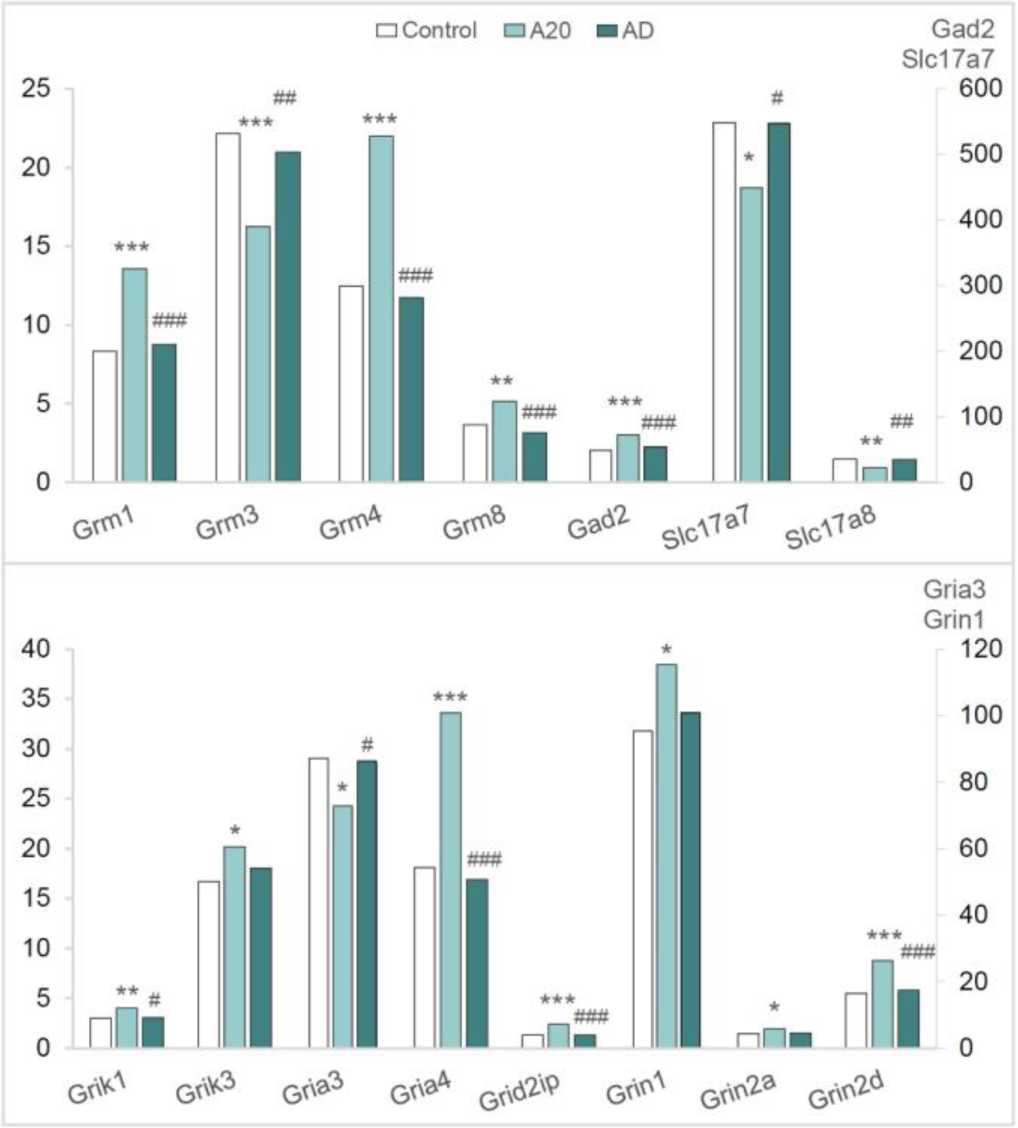
Glutamatergic-system–associated functional DEGs in the control (C) vs. A20 comparison that revert to their baseline expression after the aggression deprivation in the C vs. AD comparison in the NAcc. *C vs. A20 or C vs. AD; ^#^A20 vs. AD. Data are presented in FPKM units (fragments per kilobase of transcript per million mapped reads); C: male mice without consecutive experience of agonistic interactions; A20: male mice with consecutive 20 days of wins in daily agonistic interactions; AD: A20 male mice after the period of fighting deprivation for 14 days. \**p* < 0.05; \*\**p* < 0.01; *** *p* < 0.001; ^#^*p* < 0.05; ^###^*p* < 0.001. Additional information is shown in Supplementary Table S1.

After the period of fighting deprivation in the C vs. A20 comparison, genes *Grid1*, *Grin3a*, *Grik4*, *Grik5*, *Grm5* were upregulated, and two genes (*Gad1* and *Grm2*) were downregulated. In the comparison of A20 vs. AD mice, the expression of *Grm7* differed (Figure 7, Supplementary Table S1).

**Figure 7.**
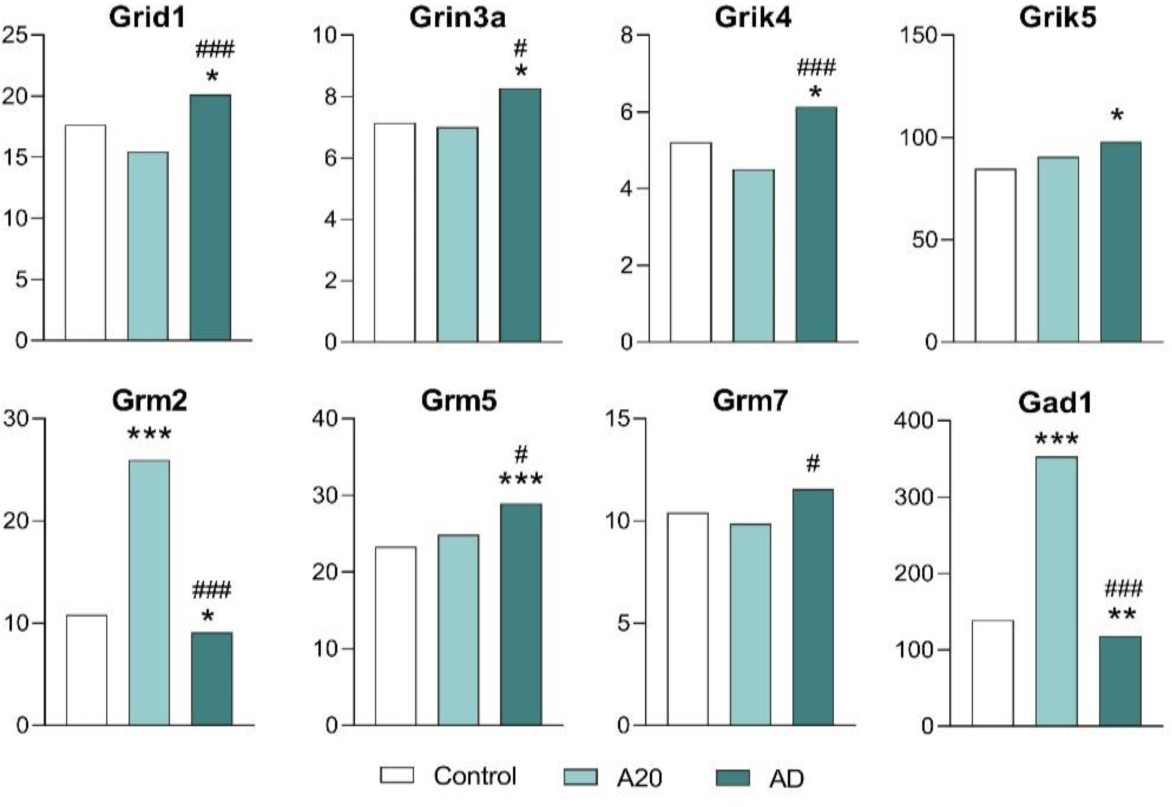
Glutamatergic-system–associated DEGs in the NAcc that are related to the observed pathology. *Control (C) vs. A20 or C vs. AD; ^#^A20 vs. AD. Data are presented in FPKM units (fragments per kilobase of transcript per million mapped reads); C: male mice without consecutive experience of agonistic interactions; A20: male mice with consecutive 20 days of wins in daily agonistic interactions; AD: A20 male mice after the period of fighting deprivation for 14 days. \**p* < 0.05; \*\**p* < 0.01; \*\*\**p* < 0.001; ^#^*p* < 0.05; ^###^*p* < 0.001. Additional information is given in Supplementary Table S1.

According to our conceptual paradigm of data analysis, glutamatergic genes *Grid1, Grin3a, Grik4, Grik5, Grm2, Grm5*, and *Grm7* (encoding ionotropic and metabotropic receptors) and the *Gad1* gene (encoding glutamate decarboxylase) may be regarded as genes associated with the observed pathology (Figure 7). Expression levels of these genes differed in the C vs. AD comparison.

Summarizing the roles both GABAergic and glutamatergic systems: all GABAergic DEGs (except for the *Gabra5a* gene), which encode GABA-A receptors (ligand-gated chloride channels), and 15 glutamatergic DEGs returned to the control expression level in AD mice after the deprivation period. Consequently, these genes cannot be regarded as participating in the development of the aggression-related pathology. By contrast, eight glutamatergic genes—*Grin3a, Grik4, Grid1, Grik5, Grm2, Grm5*, and *Grm7*— may be involved in the mechanism of relapse.

Moreover, when analyzing the role of GABAergic and glutamatergic systems, other authors [32] came to the conclusion that, for example, drug-addicted individuals can show widespread abnormalities in brain neurochemistry and function (including the systems of excitatory and inhibitory neurotransmitters glutamate and γ-aminobutyric acid - GABA, respectively, which are disturbed in addiction) that vary depending on individual characteristics (e.g., abstinence duration) and, as we propose, depending on brain regions (as demonstrated in our analyses of glutamatergic and/or GABAergic abnormalities).

### 2.3. DEGs Associated with the Observed Pathology

Within the framework of the proposed paradigm, the genes significantly differing in expression as compared with the level after fighting deprivation (C vs. AD and A20 vs. AD) can be considered genes implicated in pathologically aggressive behavior. According to the data discussed above, these include CAergic genes (*Th*, *Drd1*, *Drd2*, *Adra2c*, *Ppp1r1b*, and *Maoa*), serotonergic genes (*Htr1a*, *Htr3a*, *Htr1f*, and *Maoa*), opioidergic genes (*Oprk1*, *Pdyn*, and *Penk*), a GABAergic gene (*Gabra5*), and glutamatergic genes (*Grin3a*, *Grik4*, *Grid1*, *Grik5*, *Grm2*, *Grm5*, *Grm7*, and *Gad1*). Figure 8 presents the DEGs associated with activity of neurotransmitter systems in the NAcc; this network was constructed by means of the STRING database. DEGs *Adra2c*, *Htr1f*, *Gabra5*, *Grik4*, *Grik5*, *Grm7*, and *Htr3a* were not included in relevant highlevel interactions according to the STRING database.

**Figure 8.**
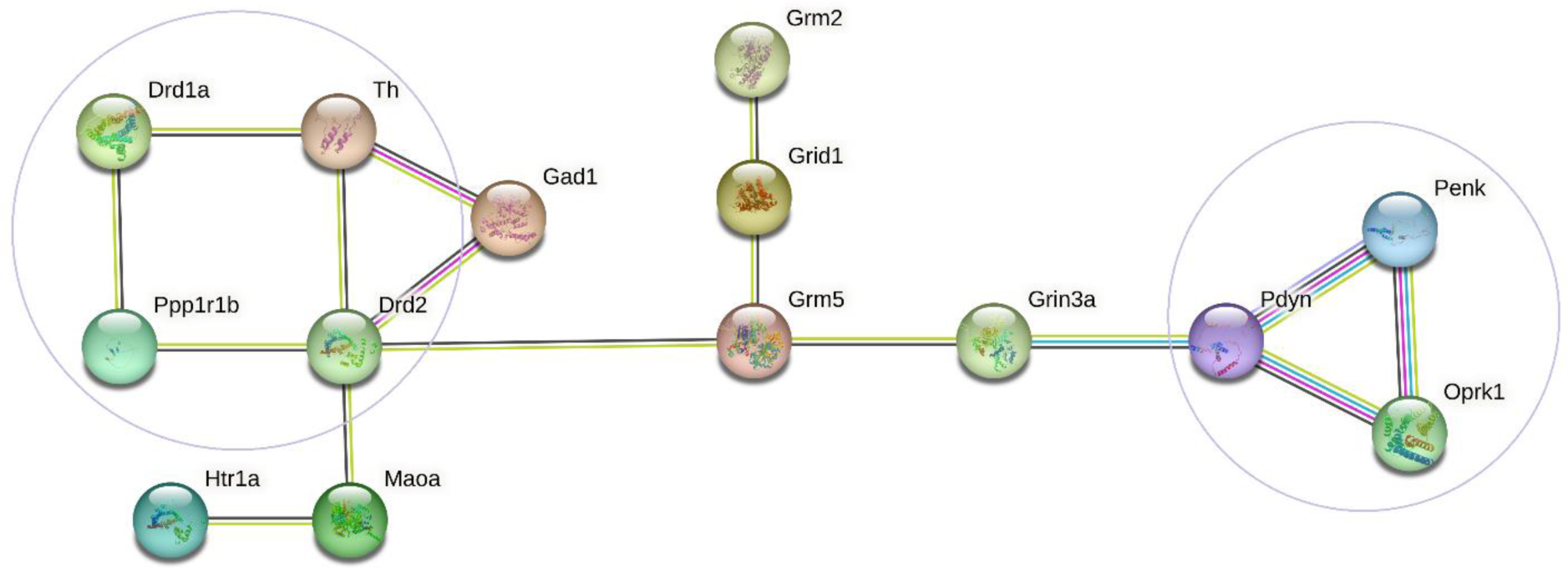
The functional enrichment network constructed by means of the STRING database (https://string-db.org/; accessed on 10 August 2023) from the DEGs associated with the activity of neurotransmitter systems in the NAcc. Lines are protein–protein associations. Purple lines indicate experimentally determined interactions; blue lines denote known interactions from curated databases; black lines show coexpression; lilac lines mean protein homology; and green lines represent results of text mining.

The STRING database (https://string-db.org/) revealed two independent relationships between the DEGs associated with СAergic systems (*Drd1*, *Drd2*, *Th*, and *PPP1r1b*) and opioidergic genes (*Oprk1*, *Pdyn*, and *Penk*), which are responsible for different processes. Nevertheless, these two groups are interconnected through glutamatergic gene *Grm5* (encoding metabotropic receptors) and glutamatergic gene *Grin3a* (encoding ionotropic receptors). This analysis suggests that the DEGs in question may contribute to the pathological addiction-like state emerging during the positive fighting experience.

Additionally, we found significant correlations between expression levels of these pathology-associated genes in control mice, in A20 mice, and in AD mice (Supplementary Table S2). A lower number of correlations between genes was found in control mice (0–6 correlations for some genes; 74 total) thus suggesting that most of these genes are independent from one another in intact mice.

The largest number of correlations between expression levels of pathology-associated genes was noticed in A20 mice (Supplementary Table S2). There were 229 significant correlations between genes: for example, 15 correlations with *Grm7* and *Htr3a*; 13 with *Grik4*, *Grm5*, *Ppp1r1b*, and *Th*; 12 with *Adra2c*; 11 with *Gad1, Grid1* and *Grm2*; and 10 with *Drd1*, *Drd2*, *Drd3*, *Oprk1*, and *Pdyn*. This finding indicates high levels of transcriptional coordination within these gene clusters in A20 mice. After the period of deprivation, we found significant correlations between pathology-related genes in AD mice in expression levels (0–9 correlations for some genes, 103 total).

In all mouse groups, correlations between CAergic (*Adra2c*, *Drd1*, *Drd2*, *Drd3*, and *PPP1r1b*) and opioidergic (*Oprk1*, *Pdyn*, and *Penk*) genes were present (Table 2). It can be theorized that these coordinated changes of gene expression are specific for the NAcc, which is responsible for reward-related mechanisms of aggression.

**Table 2.**
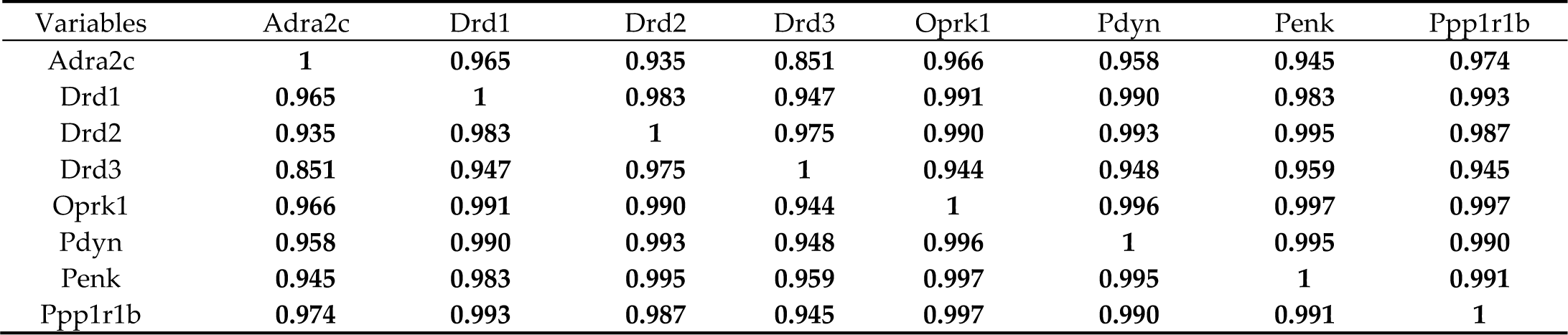
Coefficients of positive correlation between expression levels of CAergic and opioidergic genes in the NAcc of AD male mice.

| Variables | *Adra2c* | *Drd1* | *Drd2* | *Drd3* | *Oprk1* | *Pdyn* | *Penk* | *Ppp1r1b* |
| --- | --- | --- | --- | --- | --- | --- | --- | --- |
| *Adra2c* | 1 | 0.965 | 0.935 | 0.851 | 0.966 | 0.958 | 0.945 | 0.974 |
| *Drd1* | 0.965 | 1 | 0.983 | 0.947 | 0.991 | 0.990 | 0.983 | 0.993 |
| *Drd2* | 0.935 | 0.983 | 1 | 0.975 | 0.990 | 0.993 | 0.995 | 0.987 |
| *Drd3* | 0.851 | 0.947 | 0.975 | 1 | 0.944 | 0.948 | 0.959 | 0.945 |
| *Oprk1* | 0.966 | 0.991 | 0.990 | 0.944 | 1 | 0.996 | 0.997 | 0.997 |
| *Pdyn* | 0.958 | 0.990 | 0.993 | 0.948 | 0.996 | 1 | 0.995 | 0.990 |
| *Penk* | 0.945 | 0.983 | 0.995 | 0.959 | 0.997 | 0.995 | 1 | 0.991 |
| *Ppp1r1b* | 0.974 | 0.993 | 0.987 | 0.945 | 0.997 | 0.990 | 0.991 | 1 |

Of note, similar data have been obtained in human alcoholics [29]: expression of DRD1 and DRD2 strongly correlates with that of PDYN and OPRK1, indicating dysregulation of the DYN/KOR system and of DA signaling because of i) alterations in coexpression patterns of opioid genes and ii) DRD1 underexpression. This observation allowed us to hypothesize similar mechanisms of formation addictive states during an imbalance in the activity of D1 receptorand D2 receptor–containing pathways if we compare human alcoholics with our winners having the repeated experience of aggression.

## 3. Discussion

The aim of this study was to search for genes taking part in the development of a pathological condition similar to psychosis-like behavior with signs of an addiction-like state during repeated positive fighting experience in daily agonistic interactions between mice, as shown earlier in our papers [12,13,25,26]. As a criterion for the selection of genes associated with the pathology, we chose the following conceptual paradigm of data analysis: after a 14-day period of fighting deprivation (without agonistic interactions and demonstration of aggression), these genes should differ significantly in their expression compared to control mice and/or A20 mice without a return to baseline expression after the deprivation period.

In this work, we deal with the neurotransmitter genes whose products participate in CAergic, serotonergic, opioidergic, GABAergic, and glutamatergic regulatory systems in the NAcc, which, according to literature data, are involved in mechanisms of aggression accompanied by positive reinforcement and reward, as reported in experimental and psychological studies [35–37].

Summarizing the results of our study and examining our data within the proposed paradigm, a number of pathology-associated genes with changed expression in the NAcc can be considered: upregulated genes *Drd1*, *Drd2*, *Drd3*, *Adra2c*, *Ppp1r1b*, *Oprk1*, *Pdyn*, *Penk*, and *Htr1a* and downregulated genes *Th*, *Maoa*, *Htr1f*, and *Htr3a.* The GABAergic *Gabra5* gene as well as glutamatergic genes *Grin3a*, *Grik4*, *Grid1*, *Grik5*, *Grm2*, *Grm5*, *Grm7*, and *Gad1* were also included according to our above-mentioned criteria. Most of these genes significantly correlated in expression changes between one another (Supplement 1, Table S2). We assume that dopaminergic genes in the NAcc are responsible for the formation of positive emotions and development of psychosis-like behavior during prolonged activation of CAergic processes. At the same time, opioidergic genes are involved in an addiction-like state thereby promoting a relapse of aggressive behavior during withdrawal, accompanied by psychoneuropathological symptoms. According to STRING database analysis, we can say that glutamatergic proteins encoded by genes *Grm5* (coding for metabotropic receptors) and *Grin3a* (encoding ionotropic receptors) can mediate the regulation of proteins encoded by the analyzed CAergic and opioidergic genes and expression of these genes. The *Gad1* gene encoding glutamate decarboxylase is involved in the regulation of genes *Th* and *Drd2*.

We noticed a dynamic rearrangement of the functioning of genes during the formation of the observed behavioral pathology. In control animals, the number of correlations between pathology-associated genes is lower (79) than in the other groups of mice, which reflects normal functioning of genes without any external perturbations. In A20 mice, with the development of the pronounced pathology of behavior, many genes seem to participate in its development (229 genes that correlate with one another in expression). After the period of fighting deprivation, it is possible to identify the genes that retain the altered expression (only 103 genes that correlate with each other).

Previously, it has been assumed on the basis of our studies [12,25,26] that during the repeated positive fighting experience, the natural innate mechanisms regulating aggressive behavior transform into pathological ones, which are based on neurochemical shifts in the brain. It has been hypothesized that under certain circumstances, effects of endogenous opioids in chronically aggressive individuals can be reversed, and the resultant emotional and physical discomfort will eventually lead to an internal drive to aggression or to an outbreak of aggression. Thus, with prolonged aggression, neurobiological mechanisms are activated that themselves stimulate aggression. This mechanism may be another reason for the aggression relapse. Now we demonstrated that dopaminergic and opioidergic genes encoding respective proteins in the NAcc play major role in these processes.

We can also say that the psychosis-like state can be a consequence of high impulsivity developing during decreased serotonergic activity shown in many papers [12,14,61] and stimulated dopaminergic and opioidergic systems, accompanying the withdrawal after the no-fight period in the winners. Nonetheless, one should keep in mind that the development of all these states is determined by duration of positive fighting experience, and, above all, by the social context and social environment that involves the absence of punishment and of public condemnation (in humans).

The same effects can be assumed in the development of the corresponding psychopathologies in humans because escalated violence and aggression accompany the development of many psychiatric disorders: e.g., manic-depressive disorder, schizophrenia, compulsive-obsessive disorder, drug abuse, and autism [reviews: 67-69]. Recurrent aggressive behavior also takes place in professional sports, security services, and other occupations. The investigation of neurobiological mechanisms of (and factors provoking) repeated aggressive behavior is obviously an urgent task. Recent reviews on this subject [30,31,35–37,41,70] have attempted to shed light on this issue to confirm or refute previous notions about the rewarding effects of aggression attributed previously only to humans. Such animal studies have demonstrated that abnormal aggression can be excessive and serves as a reinforcement, which can lead to a pathological state during repeated positive fighting experience, in our case, to a psychosis-like state with signs of an addiction-like state.

## 4. Materials and Methods

### 4.1. Animals

Adult C57BL/6 male mice were obtained from the Animal Breeding Facility, a branch of the Institute of Bioorganic Chemistry of the RAS (Pushchino, Moscow region). Animals were housed under standard conditions (at a constant temperature of 22 ± 2 °C, under a 12:12 h light/dark regimen starting at 8:00 am, with food in pellets and water available ad libitum). Mice were weaned at 3 weeks of age and housed in groups of 8–10. The experimental cages were standard plastic cages. Experiments were performed on 10– 12-week-old animals. All procedures were in compliance with European Communities Council Directive 210/63/EU of September 22, 2010. The protocol of the study was approved by Scientific Council No. 9 of the Institute of Cytology and Genetics SB RAS of March 24, 2010, decision No. 613 (Novosibirsk).

### 4.2. Implementation of the Repeated Aggressive Experience in Male Mice

Repeated positive and negative social experiences in male mice were induced by daily agonistic interactions through the use of the sensory contact model [10], which was later renamed as the “model of chronic social conflicts” and is described in detail in some reviews [13,17]. Pairs of male mice were each placed in a cage (28 × 14 × 10 cm) bisected by a transparent perforated partition allowing the animals to see, hear, and smell each other but preventing physical contact [Figure 9].

**Figure 9.**
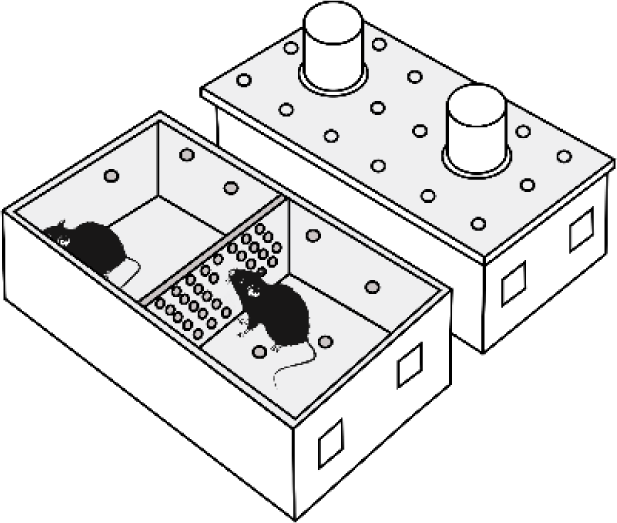
Experimental cage.

The animals were left undisturbed for 2 days to adapt to the new housing conditions and sensory contact before they were subjected to agonistic encounters. Then, every afternoon (2:00–5:00 p.m. local time) the cage cover was replaced by a transparent one, and 5 min later (the period necessary for activation), the partition was removed for 10 min to encourage agonistic interactions. The superiority of one of the mice was established within two or three encounters with the same opponent. The superior mouse would be chasing, biting, and attacking another, who would be demonstrating only defensive behavior (e.g., upright or sideways postures and withdrawal). To prevent the damage to defeated mice, the aggressive interactions between males were discontinued by lowering the partition if the strong attacking behavior lasted for 3 min (in some cases less). Each defeated mouse (loser) was exposed to the same winner for 3 days, whereas afterwards, each loser was placed after the fight into an unfamiliar cage containing an unfamiliar winning partner behind the partition. Each aggressive mouse (winner) remained in its own cage. This procedure was performed once a day for 20 days and yielded equal numbers of losers and winners. In this study, intermittent aggression was chosen for the experiment, in contrast to the strong daily aggression that was used in our previous study [16] (we distinguish three types of winners: strongly aggressive, aggressive, and intermittently aggressive). After that, the winners were deprived of the agonistic interactions for 14 days: the partition was not removed. The scheme of the experiment is presented in subsection 4.3, Figure 10.

**Figure 10.**
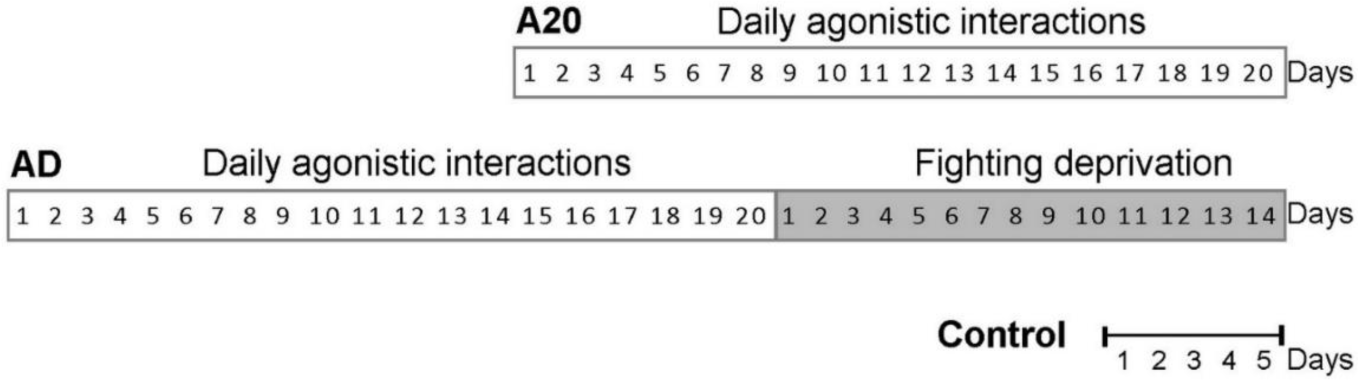
Timetable of the experiment. A20: the mice with 20-day repeated aggression accompanied by wins in the daily agonistic interactions; AD: A20 mice after 14 days of fighting deprivation; Control: mice without consecutive agonistic interactions.

### 4.3. Experiment Scheme

In this experiment, three groups of animals were analyzed: (1) controls (C), i.e., mice without a consecutive experience of agonistic interactions; (2) A20: winners, i.e., a group of mice daily intermittently aggressive during 20 days; (3) AD: fight-deprived winners after 14-day deprivation (no-fight period, without agonistic interactions), before which they were repeatedly aggressive mice during 20 days.

### 4.4. Behavioral tests for aggressive motivation and hostile behavior

Video recordings were carried out to describe the behavior of experimental animals in detail during the agonistic interactions. Among the winners (mice with positive fighting experience in 20 tests [for 20 days]), most highly aggressive animals were identified, which in daily agonistic interactions demonstrated the longest duration of aggressive motivation and hostile behaviors.

The partition test [71] was performed to estimate the aggressive motivation of mice toward the losers. The total time spent near the partition (moving near the partition, smelling and touching it with one or two paws, clutching and hanging, sticking the nose into the holes or gnawing the holes) were scored during 5 min as indices of reacting to the partner in the neighboring compartment of an experimental cage. The time during which the males showed a sideways posture or were “turning away” near the partition, was not included in the total time scored. It has previously been shown that the time spent near the partition correlates with the severity of aggressive behavior, i.e., the duration of attacks during the test.

As parameters of hostile behaviors, we analyzed the following parameters during the test when the partition was removed: total duration of the attacks, biting, and chasing a partner in agonistic interactions during 10 min; total duration of digging: here it means digging up and scattering the sawdust on the loser’s territory (kick digging or push digging the sawdust forward or backward by forepaws or hind paws); the total duration of aggressive grooming: the winner mounts the loser’s back, holds it down, and spends much time licking and nibbling at the loser’s scruff of the neck. It is regarded as a ritual form of aggression that effectively suppresses the other male mice without a physical effort, thereby replacing attacks. All these activities were aimed at inflicting physical or psychological damage on the conspecific.

The number of mice used for transcriptomic analysis was 6 for each animal group. The winners (A20) 24 h after the last agonistic interaction, fight-deprived winners (AD), and the control animals (C) were decapitated simultaneously. The NAcc was dissected by the same highly experienced experimenter according to the map presented in the Allen Mouse Brain Atlas, 2021 [https://mouse.brain-map.org/static/atlas; bregma; +1,745 - +0,745] [72]. All tissue samples were placed in an RNAlater solution (Life Technologies, Waltham, MA, USA) and were stored at −70 °C until sequencing.

### 4.5. The NAcc in the Control of Aggressive Behavior

The NAcc is a part of the reward system and plays an important role in the processing of rewarding and reinforcing stimuli (addictive drugs, sex, and exercise). According to the literature [reviews, 73,74], the NAcc is a principal target of dopaminergic neurons of the VTA, and is considered a critical brain region involved in reward and drug dependence processes [75]. Most of neurons in the NAcc are GABAergic medium spiny neurons (MSNs). The major projection neurons in this brain region are DA receptor DRD1- and DRD2-expressing neurons. Approximately 1–2% are cholinergic interneurons, and another 1–2% are GABAergic interneurons. GABAergic MSNs play an important part in the processing of reward stimuli [76], and they are regulated by DA from the VTA. Functional and molecular alterations occur within the NAcc in numerous psychoneurological pathologies, such as depression, addiction, schizophrenia, Huntington’s, Parkinson’s, and Alzheimer’s diseases, and alcoholism [74, review].

Recent research interest is concentrated also on the role of the mesolimbic dopaminergic circuit in the control of aggression reward [35–37]. It has been reported that Drd1-expressing neurons control aggression self-administration and aggression seeking in mice [70,77]. Dopaminergic projections from the VTA to the NAcc modulate DA levels [38] and aggression intensity [78,79]. Behavioral, neurochemical, neuromolecular, and neurogenomic mechanisms of prolonged experience of repeated aggression, aggression addiction, and relapse have been researched in refs. [11–13,26].

### 4.6. RNA-Seq data Analysis and Processing

The collected brain samples (control, A20, AD, n = 6 in each group) were delivered to Genoanalytica Ltd (www.genoanalytica.ru, accessed on 18 May 2023, Moscow, Russia) for RNA-Seq sequencing. The RNA-Seq method is described in details in the [24]. Briefly, total RNA was extracted using the PureLink RNA Micro Kit (Invitrogen, Waltham, Massachusetts, USA). For each tissue sample, all the extracted mRNA was used to create cDNA libraries with the help of the NEBNext mRNA Library PrepReagent Set for Illumina (New England Biolabs, Ipswich, MA USA). Sequencing of the cDNA libraries was performed on the Illumina HiSeq 2500 platform (Illumina Sequencing, San Diego, CA, USA). The target coverage was set to 20 million reads per sample on average. Cufflinks/Cuffdiff software was used to estimate gene expression levels in FPKM units (fragments per kilobase of transcript per million mapped reads) and to identify subsequently DEGs in the NAcc of male mice. Genes were considered differentially expressed at *p* < 0.05, and these data were corrected for multiple comparisons using FDR method at q < 0.05

### 4.7. The Genes That Were Analyzed in the NAcc Before and After the Deprivation Period (Supplementary Table S3)

Catecholaminergic (CAergic) systems: *Th*, *Ddc*, *Dbh*, *Маоа*, *Maob*, *Сomt*, *Slc6a2, Slc6a3, Slc18a2, Snca, Sncb, Sncg, Ppp1r1b, Drd1, Drd2, Drd3, Drd4, Drd5, Adra1a, Adra1b, Adra1d, Adra2a, Adra2b, Adra2c, Adrb1, Adrb2, Adrb3, Adrbk1,* and *Аdrbk2;*

Opioidergic and cannabinoidergic systems: *Pdyn, Penk, Pomc, Pnoc, Oprm1, Oprd1, Oprk1, Opcml, Ogfr, Ogfrl1, Cnr1, Cnr2,* and *Faah*;

Serotonergic system: *Tph2, Ddc, Maoa, Maob, Htr1a, Htr1b, Htr2a, Htr2c, Htr3a, Htr4, Htr5b, Htr6, Htr7, Htr1d, Htr1f, Htr2b, Htr3b, Htr5a, Slc6a4,* and *Slc18a2;*

GABAergic system: *Gabra1, Gabra2, Gabra3, Gabra4, Gabra5, Gabra6, Gabrb1, Gabrb2, Gabrb3, Gabrg1, Gabrg2, Gabrg3, Gabrd, Gabre, Gabrp, Gabrq, Gabbr1, Gabbr2, Gabrr1, Gabrr2, Gabrr3, Slc6a11*, and *Slc6a13*;

Glutamatergic system: *Gria1, Gria2, Gria3, Gria4, Grik1, Grik2, Grik3, Grik4, Grik5, Grin1, Grin2a, Grin2b, Grin2c, Grin2d, Grin3a, Grin3b, Grm1, Grm2, Grm3, Grm4, Grm5, Grm6, Grm7, Grm8, Grid1, Grid2, Grid2ip, Gad1, Gad2, Slc17a6, Slc17a7*, and *Slc17a8*.

Functional annotation and classification of DEGs was performed using the DAVID Bioinformatics Resources (https://david.ncifcrf.gov, ver. Dec. 2021, accessed on 20 June 20230) to describe DEGs’ functional relations and Gene Ontology terms. The Human Gene Database (http://www.genecards.org/, (accessed on 20 July 2021), Online Mendelian Inheritance in Man database (http://omim.org/, (accessed on 20 July 2021), and a human disease database (MalaCards, http://www.malacards.org, (accessed on 20 July 2021) were employed for the description and analysis of the data obtained.

## 5. Conclusions

Previously, it has been demonstrated that accumulation of effects of positive fighting experience day to day is accompanied by significant changes in social and individual behaviors and by multiple longterm changes in neurotransmitters’ synthesis, catabolism, and receptors sensitivity and in the expression of neurotransmitter-related genes in the brain, consistent with the development of a psychosis-like state with signs of an addiction-like state in mice [12,13]. After a no-fight period (aggression deprivation), the changes in expression of numerous genes and enhanced aggressiveness persist for a long period, and this phenomenon is one of the reasons for the recurrent aggression (relapse) demonstrated in our experiments [13,25,26]. Now, we have evidence that numerous genes-associated with major neurotransmitter systems (especially the opioidergic and CAergic systems) in the reward-related brain regions, in particular in the NAcc of mice—retain their altered expression after 2 weeks of the no-fight period (without agonistic interactions). In light of our proposed conceptual paradigm of data analysis, the revealed DEGs can be considered genes associated with the observed pathology that play a major role in consequences of the repeated positive fighting experience.

## 6. Limitations

In our previous publication [16], we wrote that the expression of the same genes can increase in some brain regions and diminish in others. This likely means that gene expression can depend on the function of the brain region and on the cellular environment. This state of affairs creates difficulties with finding the main target for drugs to cure the addictive state or associated pathological consequences of repeated experience of aggression.

The second problem is a pattern of changes in gene expression depending on the duration of agonistic interactions. These dynamics can differ among (and be specific for) genes, brain regions, and time points of analysis after exposure to an experimental factor, in our model, agonistic interactions. At any given moment, we can observe a certain pattern of changes and relations of genes, as estimated by means of a gene expression change or its absence.

Of great importance is also the level of aggressiveness, which can vary among winning mice. Mice can be categorized into highly aggressive mice, aggressive mice, and intermittently aggressive mice. They demonstrated aggression toward partners, however, the severity of aggressiveness could vary. It is possible that this may affect the expression of genes associated with aggression and addictive-like or other pathological behaviors. Nevertheless, it should be emphasized that the main symptoms of the development of the behavioral pathology—an increase in aggressiveness after the period of deprivation as well as high impulsivity, enhanced hyperactivity, and aggressive motivation toward partners—were well reproducible in all our experiments.

Further research is needed to elucidate the pattern of changes and possible relations between the severity of aggression and the expression of all genes associated with neurotransmitter systems in relevant brain regions in order to understand the mechanisms of action of endogenous opioidergic and СAergic genes in the neurotranscriptomic mechanisms of the addictive state.

## Abbreviations

C: control
A20: groups of intermittently daily aggressive mice during 20 days
AD: A20 mice after 14-day deprivation (no-fight period, without agonistic interactions)
5-HT: serotonin
5-HIAA: 5-hydroxyindolacetic acid
DA: dopamine
DOPAC: 3,4-dihydroxyphenyleacetic acid
DEGs: differentially expressed genes
FPKM: fragments per kilobase of transcript per million mapped reads
CAergic: catecholaminergic
GABA: γ-aminobutyric acid
MRN: midbrain raphe nuclei
MSNs: medium spiny neurons
NAcc: nucleus accumbens
STR: dorsal striatum
VTA: ventral tegmental area

## Supplementary Materials

The following supporting information can be downloaded at: www.mdpi.com/xxx/s1, Supplementary Table S1: FPKM values for neurotransmitter DEGs in the NAcc; Table S2: Correlations between expression of genes of pathology; Table S3: Neurotransmitter genes analyzed in the NAcc before and after deprivation period.

## Author Contributions

Conceptualization, N.N.K.; Data curation, D.A.S., I.L.K. and A.G.G.; Formal analysis, N.N.K., D.A.S., O.E.R., and V.N.B.; Funding acquisition, N.N.K.; Investigation D.A.S, O.E.R., V.N.B., I.L.K., A.G.G., and N.N.K.; Methodology and Supervision, N.N.K.; Writing—original draft O.E.R., N.N.K. and D.A.S.; Writing—review and editing, D.A.S., O.E.R., and V.N.B.; All authors have read and agreed to the published version of the manuscript.

## Funding

The study was supported by the Russian Science Foundation (grant No. 19-15-00026).

## Institutional Review Board Statement

All procedures were conducted in compliance with European Communities Council Directive 210/63/EU of 22 September 2010. The study protocol was approved by Scientific Council No. 9 of the Institute of Cytology and Genetics SB RAS of 24 March 2010, No. 613 (Novosibirsk, Russia).

## Informed Consent Statement

Not applicable.

## Data Availability Statement

All relevant data are available in Supplementary Materials and from the authors.

## Acknowledgments

The authors are grateful to Genoanalytica Ltd (www.genoanalytica.ru, Moscow, Russia) for conducting the technological part of the experiment and the initial statistical analysis. The English language was corrected and certified by shevchuk-editing.com.

## Conflicts of Interest

The authors declare no conflict of interest.

## Notes

### Competing Interest Statement

The authors have declared no competing interest.

## References

1. Scott, J.P. Agonistic behavior of mice and rats: A review. Am. Zool. 1966, 6, 683–701. 10.1093/icb/6.4.683.

2. Scott, J.P. Theoretical issues concerning the origin and causes of fighting. In The Physiology of Aggression and Defeat; Eleftheriou, B.E., Scott, J.P., Eds.; Plenum New-York: New York, NY, USA, 1971; pp. 11–42.

3. Brain, P. F. The adaptiveness of house mouse aggression. In P.F. Brain, D. Mainardi, S. Parmigiani (Eds.). House mouse aggression. A model for understanding the evolution of social behavior. Chur: Harwood Academic Publishers, 1979, pp. 1–21.

4. Parmigiani, S.; Brain, P. F. Effects of residence, aggressive experience and intruder familiarity on attack shown by male mice. Behav. Proc. 1983, 8, 45–47.

5. Andrade, M. L.; Kamal, K. B. H.; Brain, P. F. Effects of positive and negative fighting experience on behaviour in adult male mice. In P.F. Brain, D. Mainardi, S. Parmigiani (Eds.). House mouse aggression. A model for understanding the evolution of social behavior 1987, pp. 223–232. Harwood Academic Publishers GmbH

6. Benton, D.; Brain, P.F. The role of opioid mechanisms in social interaction and attachment. In Endorphins, Opiates and Behavioural Processes; Rodgers, R.J., Cooper, S.J., Eds.; John Wiley and Sons Ltd: London, UK, 1988; pp. 217–235.

7. Brain, P.F.; Kamal, K.B.H. Effects of prior social experience on individual aggressiveness in laboratory rodents. Rassegna Psicol. 1989, 6, 37–43.

8. Caramaschi, D.; de Boer, S.F.; de Vries, H.; Koolhaas, J.M. Development of violence in mice through repeated victory along with changes in prefrontal cortex neurochemistry. Behav. Brain Res. 2008, 189, 263–272.

9. Coppens, C. M.; de Boer, S. F.; Buwalda, B.; Koolhaas, J. M. Aggression and aspects of impulsivity in wild-type rats. Aggres. Behav. 2014. 40(4), 300–308. 10.1002/ab.21527.

10. Kudryavtseva, N.N. The sensory contact model for the study of aggressive and submissive behaviors in male mice. Aggress. Behav. 1991, 17, 285–291.

11. Kudryavtseva, N. N. An experimental approach to the study of learned aggression. Aggress. Behav. 2000, 26(3), 241–256.

12. Kudryavtseva, N.N. Psychopathology of repeated aggression: A neurobiological aspect. In Perspectives on the Psychology of Aggression, Morgan, J.P., Ed.; NOVA Science Publishers, Inc: New York, NY, USA, 2006; pp. 35–64.

13. Kudryavtseva, N.N. Positive fighting experience, addiction-like state, and relapse: Retrospective analysis of experimental studies. Aggress. Viol. Behav. 2020, 52, 101403.

14. Smagin, D.A.; Boyarskikh, U.A.; Bondar, N.P.; Filipenko, M.L.; Kudryavtseva, N.N. Reduction of serotonergic gene expression in the midbrain raphe nuclei under positive fighting experience. Adv. Biosci. Biotechnol. 2013, 4, 36–44. 10.4236/abb.2013.410A3005.

15. Smagin, D.A.; Park, J.-H.; Michurina, T.V.; Peunova, N.; Glass, Z.; Sayed, K.; Enikolopov, G. Altered hippocampal neurogenesis and amygdalar neuronal activity in adult mice with repeated experience of aggression. Front. Neurosci., 2015, 9, 443. 10.3389/fnins.2015.00443.

16. Smagin, D.A.; Galyamina, A.G.; Kovalenko, I.L.; Kudryavtseva, N.N. Altered еxpression of genes associated with major neurotransmitter systems in the reward-related brain regions of mice with positive fighting experience. Int. J. Mol. Sci. 2022, 23, 13644. 10.3390/ijms232113644

17. Kudryavtseva, N.N.; Smagin, D.A.; Kovalenko, I.L.; Vishnivetskaya, G.B. Repeated positive fighting experience in male inbred mice. Nat. Protoc. 2014, 9, 2705–2717. 10.1038/nprot.2014.156.

18. Kudryavtseva, N.N.; Kovalenko, I.L.; Smagin, D.A.; Galyamina, A.G.; Babenko, V.N. Abnormal social behaviors and dysfunction of autism-related genes associated with daily agonistic interactions in mice. In Molecular-Genetic and Statistical Techniques for Behavioral and Neural Research; Gerlai, R.T., Ed.; Academic Press: San Diego, CA, USA, 2018; pp. 309–344.

19. Babenko, V.N.; Smagin, D.A.; Kovalenko, I.L.; Galyamina, A.G.; Kudryavtseva, N.N. Differentially expressed genes of the *Slc6a* family as markers of altered brain neurotransmitter system function in pathological states in mice. Neurosci. Behav. Physiol. 2020, 50, 199–209. 10.1007/s11055-019-00888-9.

20. Babenko, V.; Redina, O.; Smagin, D.; Kovalenko, I.; Galyamina, A.; Babenko, R.; Kudryavtseva, N. Dorsal striatum transcriptome profile profound shift in repeated aggression mouse model converged to networks of 12 transcription factors after fighting deprivation. Genes 2021, 13, 21. 10.3390/genes13010021.

21. Babenko V.; Redina, O.; Smagin D.; Kovalenko I.; Galyamina A.; Kudryavtseva N. Elucidation of the landscape of alternatively spliced genes and features in the dorsal striatum of aggressive/aggression-deprived mice in the model of chronic social conflicts. Genes 2023, 14(3), 599; 10.3390/genes14030599

22. Redina, O.; Babenko, V.; Smagin, D.; Kovalenko, I.; Galyamina, A.; Efimov, V.; Kudryavtseva, N. Gene expression changes in the ventral tegmental area of male mice with alternative social behavior experience in chronic agonistic interactions. Int. J. Mol. Sci. 2020, 21, 6599. 10.3390/ijms21186599.

23. Redina, O.E.; Babenko, V.N.; Smagin, D.A.; Kovalenko, I.L.; Galyamina, A.G.; Kudryavtseva, N.N. Correlation of expression changes between genes controlling 5-HT synthesis and genes Crh and Trh in the midbrain raphe nuclei of chronically aggressive and defeated male mice. Genes 2021, 12, 1811.

24. Redina, O.E.; Babenko, V.N.; Smagin, D.A.; Kovalenko, I.L.; Galyamina, A.G.; Efimov, V.M.; Kudryavtseva, N.N. Effects of positive fighting experience and its subsequent deprivation on the expression profile of mouse hippocampal genes associated with neurogenesis. Int. J. Mol. Sci. 2023, 24(3), 3040. doi: 10.3390/ijms24033040

25. Kudryavtseva, N.N. Straub tail, the deprivation effect and addiction to aggression. In Motivation of Health Behavior, O’Neal, P.W., Ed.; NOVA Science Publishers: New York, NY, USA, 2007; pp. 97–110.

26. Kudryavtseva, N.N.; Smagin, D.A.; Bondar, N.P. Modeling fighting deprivation effect in mouse repeated aggression paradigm. Prog. Neuropsychopharmacol. Biol. Psychiatry 2011, 35, 1472–1478. 10.1016/j.pnpbp.2010.10.013.

27. Ibrahim, M.K.; Hassanein, N.M.A.; Ahmed, H.M.S. Psychopharmacological assessment of the sensory contact model as a possible model of mania. J. Glob. Biosci. 2016, 5, 3725–3741. J. Glob. Biosci. 2016, 5, 3725–3741.

28. Miczek, K.A.; Faccidomo, S.P.; Fish, E.W.; DeBold, J.F. Neurochemistry and molecular neurobiology of aggressive behavior. In Handbook of Neurochemistry and Molecular Neurobiology: Behavioral Neurochemistry, Neuroendocrinology and Molecular Neurobiology; Lajtha, A., Blaustein, J.D., Eds.; Springer: Berlin/Heidelberg, Germany, 2007; pp. 285–336.

29. Bazov, I.; Sarkisyan, D.; Kononenko, O.; Watanabe, H.; Yakovleva, T.; Hansson, A.C.; Sommer, W.H.; Spanagel, R.; Bakalkin, G. Dynorphin and kappa-opioid receptor dysregulation in the dopaminergic reward system of human alcoholics. Mol. Neurobiol. 2018, 55, 7049–7061. 10.1007/s12035-017-0844-4.

30. Covington, H.E., 3rd; Newman, E.L.; Leonard, M.Z.; Miczek, K.A. Translational models of adaptive and excessive fighting: An emerging role for neural circuits in pathological aggression. F1000Research 2019, 8, 963. 10.12688/f1000research.18883.1.

31. Haller, J. Studies into abnormal aggression in humans and rodents: Methodological and translational aspects. Neurosci. Biobehav. Rev. 2017, 76, 77–86. 10.1016/j.neubiorev.2017.02.022.

32. Moeller, S.J.; London, E.D.; Northoff, G. Neuroimaging markers of glutamatergic and GABAergic systems in drug addiction: Relationships to resting-state functional connectivity. Neurosci. Biobehav. Rev. 2016, 61, 35–52. 10.1016/j.neubiorev.2015.11.010.

33. Rodriguez-Arias, M.; Navarrete, F.; Daza-Losada, M.; Navarro, D.; Aguilar, M.A.; Berbel, P.; Minarro, J.; Manzanares, J. CB1 cannabinoid receptor-mediated aggressive behavior. Neuropharmacology 2013, 75, 172–180. 10.1016/j.neuropharm.2013.07.013.

34. Filipenko, M.L.; Alekseyenko, O.V.; Beilina, A.G.; Kamynina, T.P.; Kudryavtseva, N.N. Increase of tyrosine hydroxylase and dopamine transporter mRNA levels in ventral tegmental area of male mice under influence of repeated aggression experience. Brain Res. Mol. Brain Res. 2001, 96, 77–81. 10.1016/s0169-328x(01)00270-4.

35. Flanigan, M. E.; Russo, S. J. (2019). Recent advances in the study of aggression. Neuropsychopharmacology, 44(2), 241–244. 10.1038/s41386-018-0226-2 (Review).

36. Aleyasin, H.; Flanigan, M.E.; Russo, S.J. Neurocircuitry of aggression and aggression seeking behavior: Nose poking into brain circuitry controlling aggression. Curr. Opin. Neurobiol. 2018, 49, 184–191. 10.1016/j.conb.2018.02.013.

37. Yamaguchi, T.; Lin, D. Functions of medial hypothalamic and mesolimbic dopamine circuitries in aggression. Curr. Opinion Behav. Sci. 2018, 24, 104–112. 10.1016/j.cobeha.2018.06.011.

38. Van Erp, A. M.; Miczek, K. A. Aggressive behavior, increased accumbal dopamine, and decreased cortical serotonin in rats. J. Neurosci. 2000, 20, 9320–9325.

39. Van Erp, A.M.M.; Miczek, K.A. Increased accumbal dopamine during daily alcohol consumption and subsequent aggressive behavior in rats. Psychopharmacology 2007, 191, 679–688. 10.1007/s00213-006-0637-3

40. Couppis, M.H.; Kennedy, C.H. The rewarding effect of aggression is reduced by nucleus accumbens dopamine receptor antagonism in mice. Psychopharmacology 2008, 197(3), 449–456 DOI: 10.1007/s00213-007-1054-y

41. Golden, S.A; Heshmati, M.; Flanigan, M.; Christoffel, D.J.; Guise, K.; Pfau, M.L.; Aleyasin, H.; Menard, C.; Zhang, H.; Hodes, G.E.; Bregman, D.; Khibnik, L.; Tai, J.; Rebusi, N.; Krawitz, B.; Chaudhury, D.; Walsh, J.J.; Han, M.H.; Shapiro M.L.; Russo, S.J. Basal forebrain projections to the lateral habenula modulate aggression reward. Nature. 2016, 534(7609), 688–692. doi: 10.1038/nature18601.

42. Trapnell, C.; Hendrickson, D.G.; Sauvageau, M.; Goff, L.; Rinn, J.L.; Pachter, L. Differential analysis of gene regulation at transcript resolution with RNA-seq. Nat. Biotechnol. 2013, 31, 46–53. 10.1038/nbt.2450.

43. Svenningsson, P.; Nishi, A.; Fisone, G.; Girault, J-A.; Nairn, A.C.; Greengard, P. DARPP-32: an integrator of neurotransmission. Annu. Rev. Pharmacol. Toxico.l. 2004, 44, 269–296/ doi: 10.1146/annurev.pharmtox.44.101802.121415

44. Kudriavtseva, N.N.; Bakshtanovskaia, I.V. The neurochemical control of aggression and submission. Zh. Vyssh. Nerv. Deiat. Im. I P Pavlova 1991, 41, 459–466.

45. Bondar, N.P.; Boyarskikh, U.A.; Kovalenko, I.L.; Filipenko, M.L.; Kudryavtseva, N.N. Molecular implications of repeated aggression: *Th, Dat1, Snca* and *Bdn*f gene expression in the VTA of victorious male mice. PLoS ONE 2009, 4, e4190. 10.1371/journal.pone.0004190.

46. Kudryavtseva, N.N.; Avgustinovich, D.F.; Bondar, N.P.; Tenditnik, M.V.; Kovalenko, I.L. An experimental approach for the study of psychotropic drug effects under simulated clinical conditions. Curr. Drug Met. 2008, 9(4), 352–360.

47. Kudryavtseva, N.N.; Lipina, T.V.; Koryakina, L.A. Effects of haloperidol on communicative and aggressive behavior in male mice with different experiences of aggression. Pharmacol. Biochem. Behav. 1999, 63, 229–236. 10.1016/s0091-3057(98)00227-5.

48. Bondar’, N.P.; Kudriavtseva, N.N. Effect of the D1-receptor antagonist SCH-23390 on the individual and aggressive behavior in male mice with various aggression experience. Ross. Fiziol. Zh. Im. I.M. Sechenova. 2003, 89(8), 992–1000.

49. Kudriavtseva, N.N.; Dolgov, V.V.; Avgustinovich, D.F.; Alekseenko, O.V.; Lipina, T.V.; Koriakina, L.A. Modifying effect of the repeated experience of agonistic confrontations on effect of naltrexone in male mice. Ross. Fiziol. Zh. Im. I. M. Sechenova 2001, 87, 227–238.

50. Lipina, T.V.; Avgustinovich, D.F.; Koriakina, L.A.; Alekseenko, O.V.; Kudriavtseva, N.N. Differences in the effects of naltrexone on the communicative and aggressive behaviors of subjects with different experiences of social conquests. Eksp. Klin. Farmakol. 1998, 61, 13–18.

51. Bondar, N.P.; Smagin, D.A.; Kudryavtseva, N.N. Effects of single and chronic naltrexone treatment on agonistic behavior of male mice with repeated experience of aggression. Psychopharmacol. Biol. Narcol. 2011, 11, 2688–2700.

52. Kudryavtseva, N.N.; Gerrits, M.A.; Avgustinovich, D.F.; Tenditnik, M.V.; van Ree, J.M. Modulation of anxiety-related behaviors by mu- and kappa-opioid receptor agonists depends on the social status of mice. Peptides 2004, 25, 1355–1363. 10.1016/j.peptides.2004.05.005.

53. Kudryavtseva, N.N.; Gerrits, M.A.; Avgustinovich, D.F.; Tenditnik, M.V.; Van Ree, J.M. Anxiety and ethanol consumption in victorious and defeated mice; effect of kappa-opioid receptor activation. Eur. Neuropsychopharmacol. 2006 16(7), 504–511. doi: 10.1016/j.euroneuro.2006.01.002.

54. Goloshchapov, A.V.; Filipenko, M.L; Bondar, N.P.; Kudryavtseva, N.M.; Van Ree, J.M. Decrease of kappa-opioid receptor mRNA level in ventral tegmental area of male mice after repeated experience of aggression. Brain. Res. Mol. Brain Res. 2005, 135(1-2), 290–292. doi: 10.1016/j.molbrainres.2004.11.009.

55. Stefano, G.B.; Goumon, Y.; Casares, F.; Cadet, P.; Fricchione, G. L.; Rialas, C.; Bianchi, E. Endogenous morphine. Trends Neurosci. 2000, 23(9), 436–442

56. Neri, C.; Guarna, M.; Bianchi, E.; Sonetti, D.; Matteucci, G.; Stefano, G. B. Endogenous morphine and codeine in the brain of nonhuman primate. Med. Sci. Mon.r, 2004, 10(6), MS1–5

57. Kream, R. M.; Stefano, G. B.; Ptáček, R. Psychiatric implications of endogenous morphine: Up-to-date review. Folia Biologica (Praha) 2010), 56(6), 231–241.

58. Tordjman, S.; Carlier, M.; Cohen, D.; Cesselin, F.; Bourgoin, S.; Colas-Linhart, N.; Peti et, A.; Perez-Diaz, F.; Hamon, M.; Roubertoux, P.L. Aggression and the three opioid families (endorphins, enkephalins, and dynorphins) in mice. Behav. Genet. 2003, 33, 529–536. 10.1023/a:1025774716976

59. Kulikov, A.V.; Kozlachkova, E.Y.; Kudryavtseva, N.N.; Popova, N.K. Correlation between tryptophan hydroxylase activity in the brain and predisposition to pinch-induced catalepsy in mice. Pharmacol Biochem Behav. 1995, 50(3), 431–435. doi: 10.1016/0091-3057(94)00293-r.

60. Amstislavskaya, T.G., Kudryavtseva, N.N. Effect of repeated experience of victory and defeat in daily agonistic confrontations on brain tryptophan hydroxylase activity. FEBS Lett. 1997, 406 (1–2), 106–108. doi: 10.1016/s0014-5793(97)00252-4.

61. Kudryavtseva, N.N.; Smagin, D.A.; Kovalenko, I.L.; Galyamina, A.G.; Vishnivetskaya, G.B.; Babenko, V.N.; Orlov, Y.L. Serotonergic genes in the development of anxiety/depression-like state and pathology of aggressive behavior in male mice: RNA-seq data. Mol. Biol. 2017, 51, 288–300. 10.7868/S0026898417020136.

62. Bondar’, N.P.; Kudriavtseva, N.N. Effect of buspirone on aggressive and anxiety behavior of male mice with various aggressive experience. Eksp. Klin. Farmakol. 2003 66(4), 12–16.

63. Brown, G.L.; Ebert, M.H.; Goyer, P.F.; Jimerson, D.C.; Klein, W.J.; Bunney, W.E.; Goodwin, F.K. Aggression, suicide, and serotonin: relationships to CSF amine metabolites. Am. J. Psychiatry. 1982 139, 6, 741–746. doi: 10.1176/ajp.139.6.741.

64. Coccaro E.F. Impulsive aggression and central serotonergic system function in humans: an example of a dimensional brain-behavior relationship. Int. Clin. Psychopharmacol. 1992, 7(1), 3–12 DOI: 10.1097/00004850-199200710-00001.

65. Glick, A.R. The role of serotonin in impulsive aggression, suicide, and homicide in adolescents and adults: A literature review. Int J. Adolesc. Med. Health 2015, 27, 143–150. 10.1515/ijamh-2015-5005.

66. Ferrari, P.F.; Palanza, P.; Parmigiani, S.; de Almeida, R.M.M.; Miczek, K.A. Serotonin and aggressive behavior in rodents and nonhuman primates: predispositions and plasticity. Eur. J. Pharmacol. 2005, 526(1–3), 259–273. DOI: 10.1016/j.ejphar.2005.10.002

67. de Boer, S.F. Animal models of excessive aggression: Implications for human aggression and violence. Curr. Opin. Psychol. 2018, 19, 81–87. 10.1016/j.copsyc.2017.04.006.

68. De Boer, S.F.; Buwalda, B.; Koolhaas, J.M. Untangling the neurobiology of coping styles in rodents: Towards neural mechanisms underlying individual differences in disease susceptibility. Neurosci. Biobehav. Rev. 2017, 74, 401–422. 10.1016/j.neubiorev.2016.07.008.

69. Diagnostic and Statistical Manual of Mental Disorders, Fifth Edition (DSM-5™), 2013

70. Golden, S.A.; Shaham, Y. Aggression addiction and relapse: A new frontier in psychiatry. Neuropsychopharmacology 2018, 43, 224–225. 2018, *43*, 224–225. 10.1038/npp.2017.173.

71. Kudryavtseva, N.N. Use of the “partition” test in behavioral and pharmacological experiments. Neurosci Behav Physiol. 2003, 33, 5, 461–471. doi: 10.1023/a:1023411217051. Review.

72. The Allen Mouse Brain Atlas. Available online: http://mouse.brain-map.org/static/atlas (accessed on 24 April 2021).

73. Robison, A.J.; Nestler, E.J. Transcriptional and epigenetic mechanisms of addiction. Nat. Rev. Neurosci. 2011, 12, 623–637.

74. Bayassi-Jakowicka, M.; Lietzau, G.; Czuba, E.; Patrone, C.; Kowiański, P. More than addiction -the nucleus accumbens contribution to development of mental disorders and neurodegenerative diseases. Int. J. Mol. Sci. 2022 23(5), 2618. doi: 10.3390/ijms23052618

75. Olsen, C.M. Natural rewards, neuroplasticity, and non-drug addictions. Neuropharmacology 2011, 61, 1109–1122.

76. Meredith, G.E.; Pennartz, C.M.; Groenewegen, H.J. The cellular framework for chemical signalling in the nucleus accumbens. Prog. Brain Res. 1993, 99, 3–24. 10.1016/s0079-6123(08)61335-7.

77. Golden, S.A.; Jin, M.; Heins, C.; Venniro, M.; Michaelides, M.; Shaham, Y. Nucleus accumbens Drd1-expressing neurons control aggression self-administration and aggression seeking in mice. J. Neurosci. 2019, 39, 2482–2496. 10.1523/JNEUROSCI.2409-18.2019.

78. Yu Q, Teixeira CM, Mahadevia D, Huang Y, Balsam D, Mann JJ, Gingrich JA, Ansorge MS. Dopamine and serotonin signaling during two sensitive developmental periods differentially impact adult aggressive and affective behaviors in mice. Mol. Psychiatry 2014, 6,688–98. doi: 10.1038/mp.2014.10.

79. Miczek, K.A.; Takahashi, A.; Gobrogge, K.L.; Hwa, L.S.; de Almeida, R.M. Escalated aggression in animal models: Shedding new light on mesocorticolimbic circuits. Curr. Opin. Behav. Sci. 2015, 3, 90–95. 10.1016/j.cobeha.2015.02.007.

